# Loss of MECP2 leads to activation of p53 and neuronal senescence

**DOI:** 10.1101/130401

**Authors:** M. Ohashi, D. Allen, P. Lee, E. Korsakova, K. Fu, B. Vargas, J. Cinkornpumin, C. Salas, J. Park, I. Germanguz, K. Chronis, E. Kuoy, S. Tran, G. Xiao, M. Pellegrini, K. Plath, WE. Lowry

**Author notes:** To whom correspondence should be addressed/lead contact: William Lowry and Kathrin Plath.

## Abstract

To determine the role for mutations of MECP2 in Rett Syndrome, we generated isogenic lines of human iPSCs (hiPSCs), neural progenitor cells (NPCs), and neurons from patient fibroblasts with and without MECP2 expression in an attempt to recapitulate disease phenotypes *in vitro*. Molecular profiling uncovered neuronal specific gene expression changes including induction of a Senescence Associated Secretory Phenotype (SASP) program. Patient derived Neurons made without MECP2 show signs of stress, including induction of p53, and senescence. The induction of p53 appeared to affect dendritic branching in Rett neurons, as p53 inhibition restored dendritic complexity. These disease-in-a-dish data suggest that loss of MECP2 can lead to dendritic defects due to an increase in aspects of neuronal aging.

## Introduction

Rett Syndrome is a disease associated with loss of function mutations in the gene MECP2, which was originally identified as encoding a methylated DNA binding protein^1–3^. Patient symptoms include microcephaly, intellectual disability, facial dysmorphia, and seizure activity^4,5^. Studies in murine models recapitulate many of the patient phenotypes and have recently identified a role for MECP2 particularly in inhibitory neurons^6–9^. These studies demonstrated that loss of MECP2 can lead to defects in transcription^10–12^, dendritic branching^13^, nuclear size^3^, and AKT signaling^14^.

MECP2 has also been described as a transcription factor with specific targets^10,11,13^, and more broadly as either a transcriptional activator^14^ or repressor^15–18^. However, despite decades of research on MECP2, it is still unclear how mutations in this protein lead to patient symptoms^3,14,19–21^. To confirm findings made in other models and further study these in a human system, some have turned to modeling Rett Syndrome *in vitro* by taking advantage of Disease in a dish approaches. This involves making hiPSCs from patient somatic cells, or using genome engineering to introduce mutations into WT human pluripotent stem cells. In either case, the pluripotent stem cells created are then differentiated toward the neural lineage, and then comparisons can be made between cells that express MECP2 or lack it.

Some of these studies have even taken advantage of isogenic or congenic cells lines to identify both transcriptional and electrophysiological effects of loss of MECP2 in human in vitro models^14,22^. In the current study, we also sought to mitigate the effect of genetic background and variability of differentiation by taking advantage of several congenic lines of hiPSCs that either express the WT allele or the mutant allele leading to cells that express or lack MECP2^23^. This allowed for detailed molecular analyses of hiPSCs, NPCs and neurons with and without MECP2 under the same genetic background. In addition, several lines were made and analyzed in each category to avoid variance in differentiation potential amongst isogenic lines. Furthermore, congenic lines were made from two patients with different mutations to highlight only those phenotypes associated with loss of MECP2 expression and not genetic background or variance in hPSC differentiation. Finally, we validated many of these findings using siRNA silencing of MECP2 in WT cells of a distinct genetic background. In comparing neurons from Rett patients as well as those with MECP2 silenced by siRNA, it is clear that loss of MECP2 leads to induction of p53 and senescence, potentially opening a new avenue of investigation for this intellectual disability syndrome.

## Results

### A congenic model of Rett Syndrome *in vitro*

To determine how loss of MECP2 expression leads to defects in the nervous system we generated a disease-in-a-dish model using iPSCs. Cognizant of the fact that differentiation from hPSCs is highly variable across individual lines, culture conditions, and time, we developed an isogenic model to study Rett Syndrome *in vitro* to remove the confound of genetic background^23^. Because female patients with Rett Syndrome are usually heterozygous for mutant alleles of *MECP2*, fibroblasts isolated from these patients display a mosaic pattern where roughly half the cells express either the mutant or WT allele. This is shown in Figure 1A, where fibroblasts isolated from two patients with distinct mutant alleles of MECP2 (R982 and R567) showed that roughly half the cells express MECP2 while the other half lacked detectable amounts of this protein. One of these mutant alleles is predicted to lead to a premature stop codon, while the other leads to failed transcriptional termination. Reprogramming to iPSCs using a small set of transcription factors has been shown to happen at the clonal level, such that individual reprogramming events in single fibroblasts generate isolated hiPSC clones^24^. Therefore, reprogramming of mosaic fibroblast cultures from two different patients generated single hiPSC clones that either expressed MECP2 protein or lacked it (Fig 1B) (Method described in a previous study^25^). In addition, our work and that of others has shown that under standard conditions, the inactive X chromosome in human fibroblasts does not reactivate upon reprogramming to the pluripotent state^23,25,26^, which is distinct from murine reprogramming^27^.

**Figure 1.**
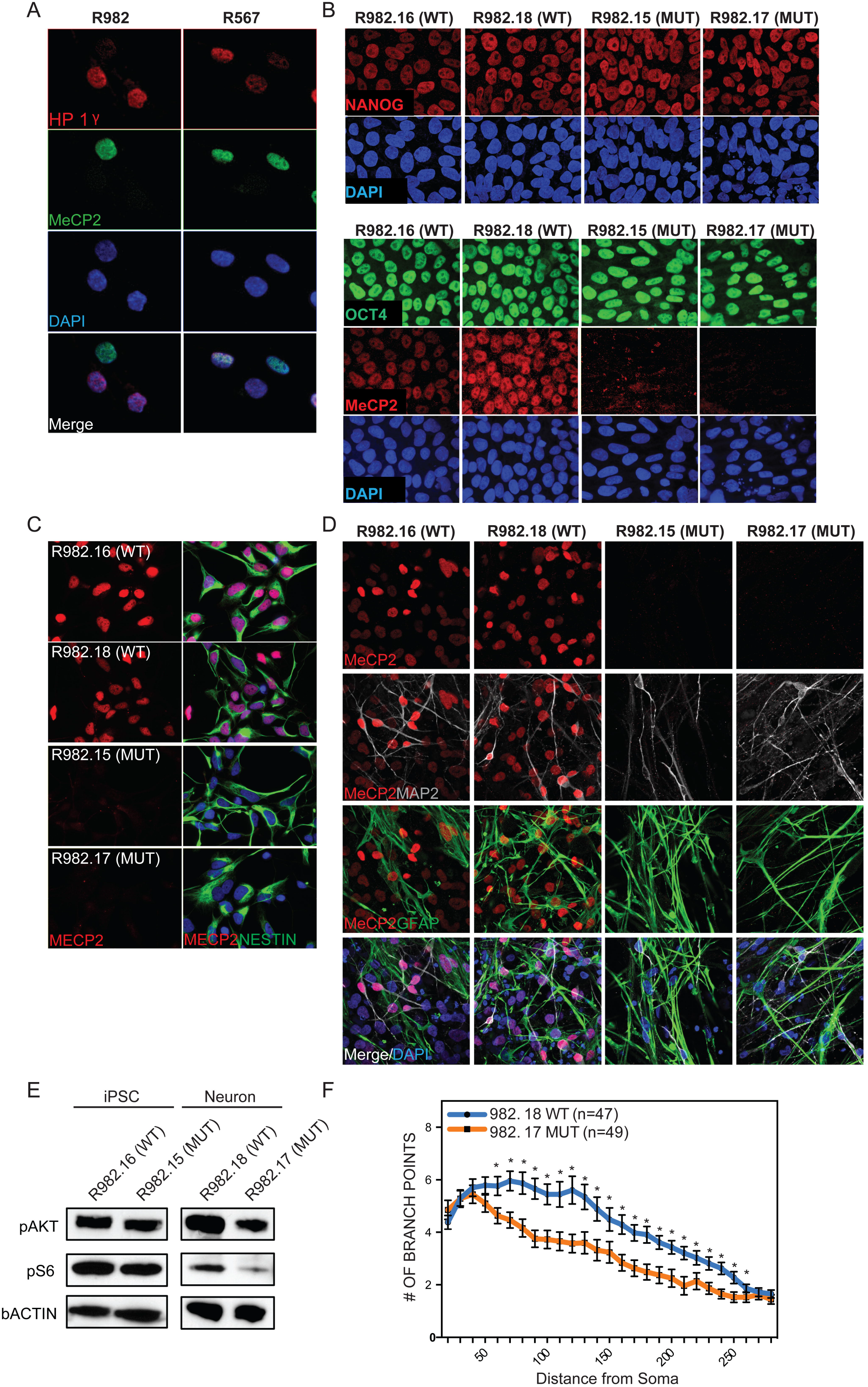
Generation of isogenic model of Rett Syndrome *in vitro*. **A**, Fibroblasts isolated from Rett Syndrome patients (R982 and R567) heterozygous for MECP2 mutations exhibit a mosaic pattern of MECP2 expression due to random XCI. Note that roughly 50% of fibroblasts from each patient express MECP2. **B**, Multiple isogenic hiPSC lines were produced from patient 982 with a typical Yamanaka protocol yielding individual isogenic clones with and without MECP2 expression from the same patient, as judged by NANOG and OCT4 staining. **C**, Specification of 982 patient derived hiPSCs towards neural progenitor cells yielded homogenous cultures of NPCs with and without MECP2. **D**, terminal differentiation of 982 patient derived NPCs towards neurons and glial by growth factor withdrawal yielded normal neural derivatives as measured by immunostaining for MAP2 and GFAP. **E**, MECP2+ and MECP2- hiPSCs and neurons were generated from patient 982 (R982.16 and R982.15) and assayed for activity of the AKT pathway by western blot with antibodies that recognize the active forms of Akt and its downstream target S6. **F**, Sholl assay of dendritic complexity was performed on WT vs MUT neurons derived from patient 982. Increased # of branch points indicates increased dendritic complexity, measured as a function of distance from the cell body. *p value < 0.05 according to student’s t test. Bar graphs represent mean +/− SEM.

Thus, we were able to create multiple lines of hiPSCs with and without MECP2 from individual patients and thereby control for differences in genetic background (shown in Fig 1B are clones made from patient 982, clones from 567 look similar). The hiPSCs generated from fibroblasts of both patients appeared to be unaffected by the lack of MECP2, expressed all appropriate markers, and successfully generated teratomas upon injection into the testes of immunocompromised mice, consistent with previous hiPSC models for loss of MECP2 (Fig S1A)^14,28–30^. Lack of MECP2 in patient-derived cells and specificity of antibody was also confirmed by western blot (Fig S1B). Importantly, we never observed reactivation of the silenced X chromosome that would have resulted in re-expression of the WT allele of MECP2 in any cultures regardless of differentiation status or passage. This is consistent with previous data showing that despite evidence for erosion of isolated portions of the silenced X chromosome^31^, the portion containing the MECP2 locus was not affected by reprogramming or differentiation.

As Rett Syndrome primarily afflicts the nervous system and MECP2 is most highly expressed in neurons, we first generated neural progenitor cells from all of the hiPSCs lines following standard protocols^32^. Across at least two lines per patient with and without MECP2, we measured the rate of neuralization, the morphology of NPCs, and expression of typical marker genes. We were unable to detect consistent differences in these properties between multiple clones of both WT and MECP2-lines derived from both patients (Fig 1C and Fig S1C). Furthermore, the growth rate of NPCs with and without MECP2 was not consistently different in NPCs made from either patient (Fig S1D). Next, the NPCs were further differentiated by a non-directed differentiation approach that yields both neurons and glia (growth factor withdrawal^33^) (Fig 1D). All NPCs from both patients produced neurons and glia at the same rate (Fig S1E and F).

Previous studies have also shown that loss of MECP2 in neurons can lead to a decrease in AKT signaling^14^. A similar pattern was observed here in mutant neurons generated from Rett patient hiPSCs as measured by phosphorylation of AKT and S6, while hiPSCs themselves did not seem to be affected by loss of MECP2 (Fig 1E). Dendritic complexity has been shown extensively to be reliant on MECP2 expression in various models of Rett Syndrome, and we found a statistically significant decrease in complexity in neurons made in the absence of MECP2 by Sholl assay (Fig 1F). In addition, we observed qualitative differences in basic neuronal morphology between WT and mutant neurons, where the neurons lacking MECP2 had shorter, thicker processes, and their soma was not as well defined.

### Loss of MECP2 affects the transcriptome of neurons

It has been suggested that loss of MECP2 only affects gene expression in neurons as opposed to the hPSCs and NPCs from which they were derived^14^. We sought to determine whether gene expression was affected in hiPSCs, NPCs or neurons in this patient derived *in vitro* model. To optimize the search for molecular effects of loss of MECP2 in hiPSCs, NPCs or neurons, we used defined neuronal cultures by following the newly established 3i (three inhibitor) method to create populations of human interneurons (Fig 2A)^34^. Interneurons are particularly relevant in the study of Rett Syndrome as interneuron-specific deletion of Mecp2 in mice recapitulates many of the disease symptoms^6,8,35,36^. We validated the purity and quality of differentiation at each step by immunostaining for markers typical of particular cell types (SOX2, SOX1 and NESTIN as well as FOXG1 and NKX2.1 for NPCs; and Tuj1, MAP2 and GABA for interneurons) in both WT and MUT cultures followed by quantification (Fig S2A and B). We first assessed whether interneurons lacking MECP2 also showed diminished dendritic branching. In fact, in patient-derived interneurons made by 3i, defects in dendritic branching as measured by the number of endpoints were clearly observed (Fig 2A).

**Figure 2.**
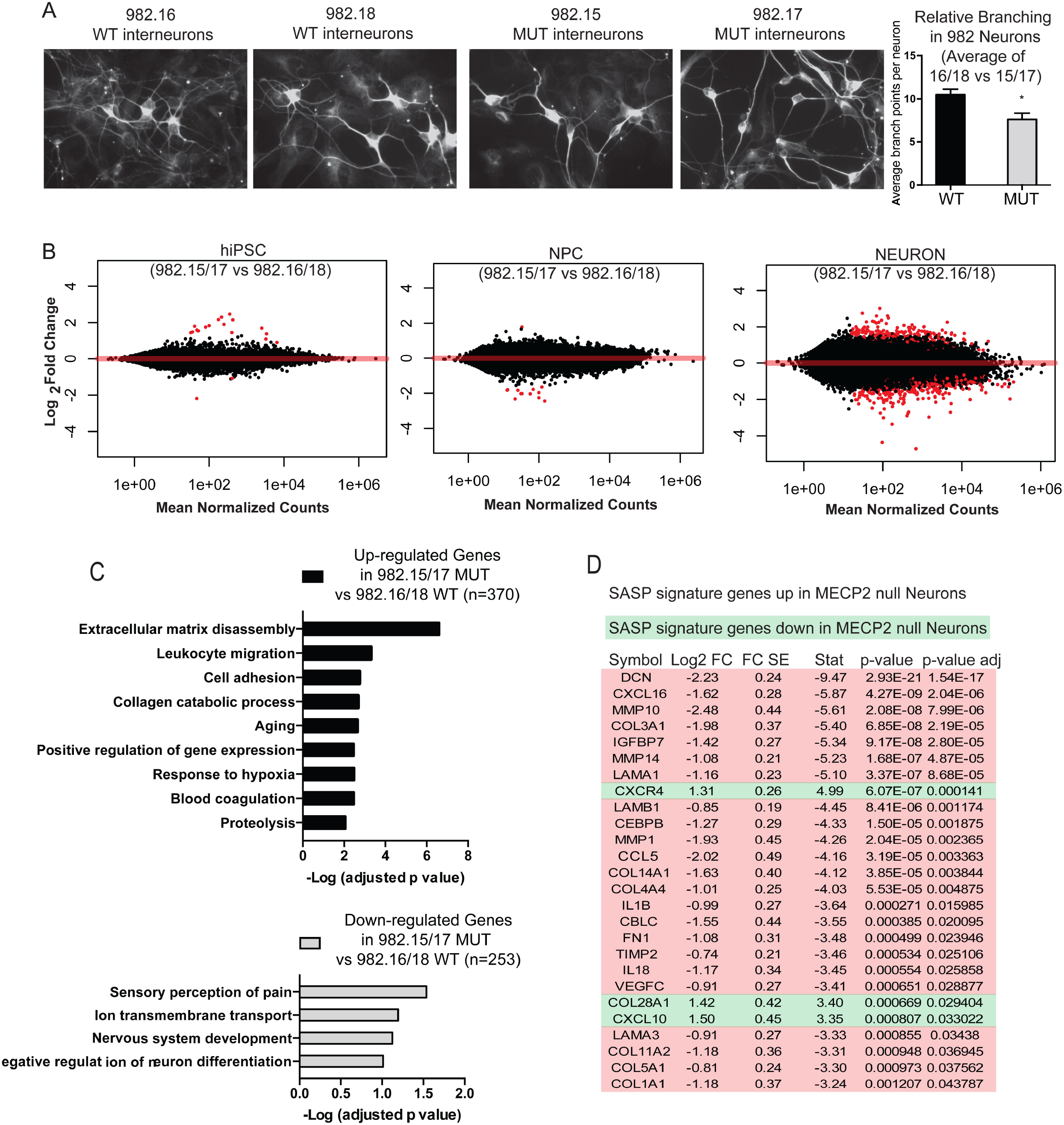
Loss of MECP2 is associated with differential gene expression in neurons. **A**, Immunostaining neurons generated from patient 982 for TuJ1, a neuronal-specific marker. **Right**, quantification of dendritic complexity by counting endpoints shows a significant difference between neurons with and without MECP2 made from patient 982. **B**, Volcano plots of differentially expressed genes (DEGs) in hiPSCs, NPCs and Neurons shows that loss of MECP2 has a profound effect on gene expression in neurons. **C**, Gene ontological analysis of DEGs increased versus decreased in MECP2 null neurons. **D**, An examination of SASP genes in neurons. A high proportion of SASP signature genes were upregulated in MECP2 null neurons versus WT neurons.

We therefore proceeded with deep RNA seq (>120 million reads per sample) of hiPSC, NPC and interneuron cultures. With such sequencing depth, it was possible to analyze the RNA-seq reads for the known mutations present in the patients from which these lines were made (Fig S2C). This analysis demonstrated that each line studied expressed strictly either the WT or mutant allele of MECP2, and that XCI status was unchanged even after extensive differentiation to neurons.

We quantified the expression level of MECP2 in WT cells across these three stages of development and found that the average RPKM was 3.1 for hiPSCs, 4.3 for NPCs, and 7.75 for interneuron cultures. This is consistent with consensus that MECP2 is enriched in neuronal cells, but also demonstrates that it could potentially be relevant to hiPSC and NPC physiology as well. However, high stringency analyses (FDR <0.05) of the RNA-seq data yielded very few gene expression changes due to loss of MECP2 in hiPSCs or NPCs derived from Rett patients (Fig 2B), consistent with Li et al^14^. On the other hand, interneuron cultures made from patient 982 showed many gene expression changes when comparing two individual WT and MUT clones (Fig 2B). Gene ontology analysis uncovered many neuronal physiology-related pathways were downregulated due to loss of MECP2 in neurons, while genes associated with extracellular remodeling and cell migration appeared to be induced (Fig 2C).

Probing RNA-seq data, we also found that MECP2 null interneuron cultures showed a strong increase in a group of genes that are known to be induced by senescent cells, known as the Senescence Associate Secretory Program (SASP). The vast majority of SASP genes that were changed in MECP2 null neurons were upregulated as opposed to downregulated, suggesting a strong pattern of SASP induction (Fig 2D). The only previous report linking MECP2 loss to senescence was performed by partial silencing of this protein in mesenchymal stem cells, but the results were consistent with those shown here for patient derived MECP2 null fibroblasts^37^. The induction of SASP was intriguing in light of the fact that, while attempting to make clones of fibroblasts from patients with Rett syndrome, we repeatedly found that clones lacking MECP2 did not expand well after passage (14 MECP2 null clones were created, none expanded), while clones expressing the WT allele expanded without problem (42 MECP2+ clones were created, and 4 out of 4 all expanded).

To determine whether MECP2 null fibroblasts encounter senescence, we performed an assay to detect endogenous beta-galactosidase, which is known to be a hallmark of this process^38^. Indeed, MECP2 null fibroblasts showed strong activity in this senescence assay (Fig 3A). We did not encounter such difficulties with clonal expansion once hiPSCs or hiPSC-derived NPCs were made from patients, presumably because during reprogramming, telomerase is strongly induced to restore telomere length at least beyond the critical threshold^39–44^. In fact, our RNA-seq data showed that hiPSCs made from patients had very high expression of TERT, and NPCs still expressed moderate levels, while neurons did not express appreciable levels (average RPKM for TERT: hiPSC, 8.8; NPC, 1.6; neuron, 0.006). Importantly, the same endogenous galactosidase activity assay on interneurons showed a dramatic increase in senescence activity in neurons lacking MECP2 (Fig 3B). These data indicate that loss of MECP2 leads to not only induction of SASP, but also a *bona fide* senescence program in neurons.

**Figure 3.**
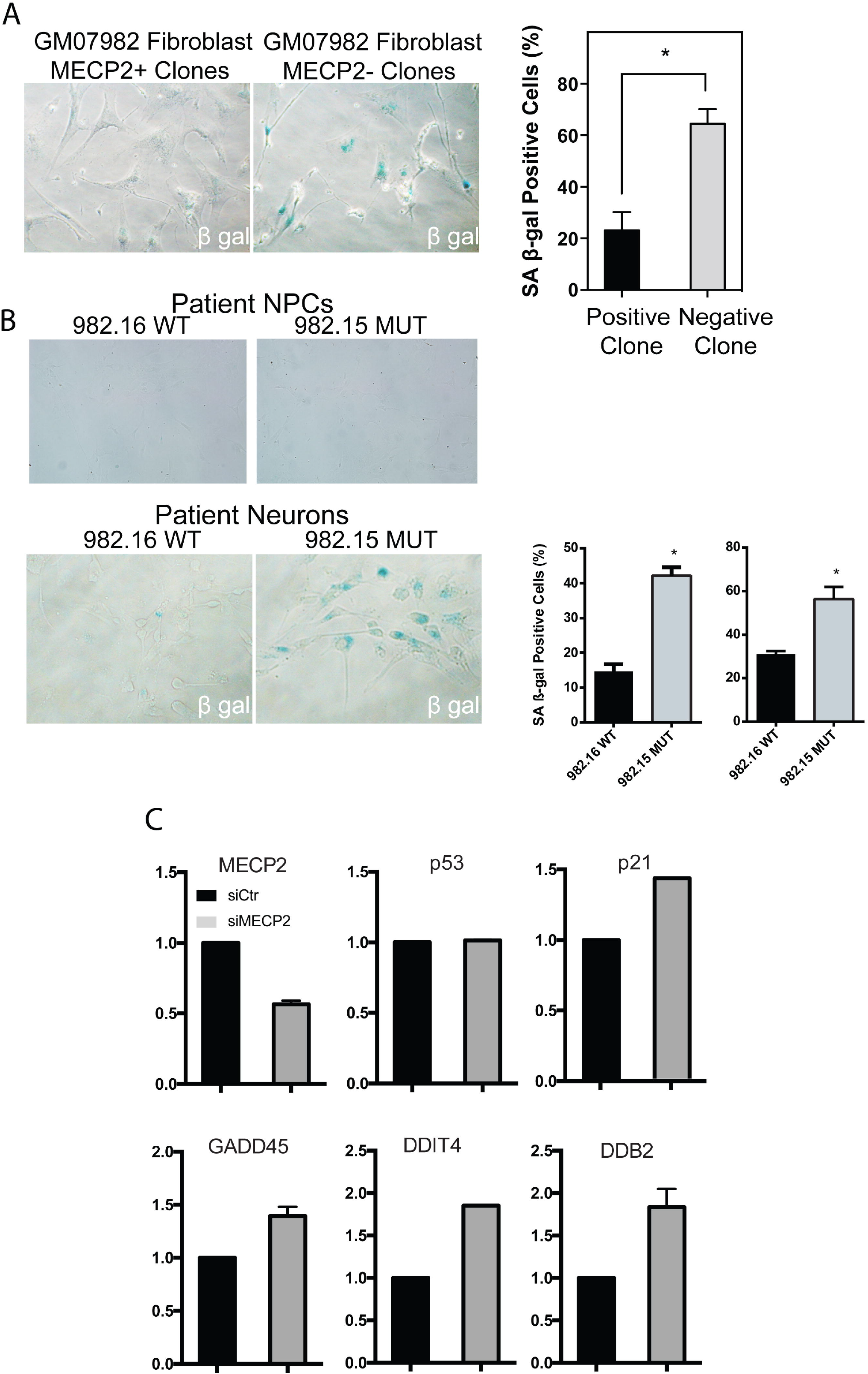
Loss of MECP2 leads to senescence and signs of telomere dysfunction. **A**, Cells undergoing senescence show upregulation of endogenous b-galactosidase activity. Patient skin-derived clones of fibroblasts lacking MECP2 showed strong b-gal activity, while those of WT fibroblasts did not. **B**, TOP, The senescence assay applied to neural progenitors derived from Rett patients did not show significant senescence activity. BOTTOM, patient derived neuronal cultures showed a strong increase in the absence of MECP2 (quantification across independent lines shown on right). **C**, RT-PCR for p53 targets after siRNA treatment of WT neurons.

### Induction of P53 in the absence of MECP2

Cellular senescence programs are known to be regulated by p53, which can then activate various response pathways downstream such as DNA repair and apoptosis^52^. Interestingly, p53 induction due to telomere shortening was previously shown to cause defects in dendritic branching^53,54^, which is also the dominant phenotype in Rett Syndrome. To begin to look for hallmarks of p53 induction in the absence of MECP2, we performed RT-PCR for p53 related targets in cells with silencing of MECP2 by siRNA (Fig S3). This assay suggested that decreased MECP2 levels led to induction of p53 related target genes such as p21, GADD45, DDIT4, and DDB2 (Fig 3C).

To determine the effect of loss of MECP2 in relation to cell stress pathways at the protein level, we performed immunostaining for H2aX, PML, p53, and p21 in neurons with and without MECP2. Staining for each of these markers showed strong increases in expression/levels of these markers of cell stress in patient-derived NPCs, neurons, and also after silencing of MECP2 in both NPCs and neurons (Fig 4A-D). WT NPCs with silencing of MECP2 by siRNA and neurons lacking MECP2 also showed clear induction of these marks.

**Figure 4.**
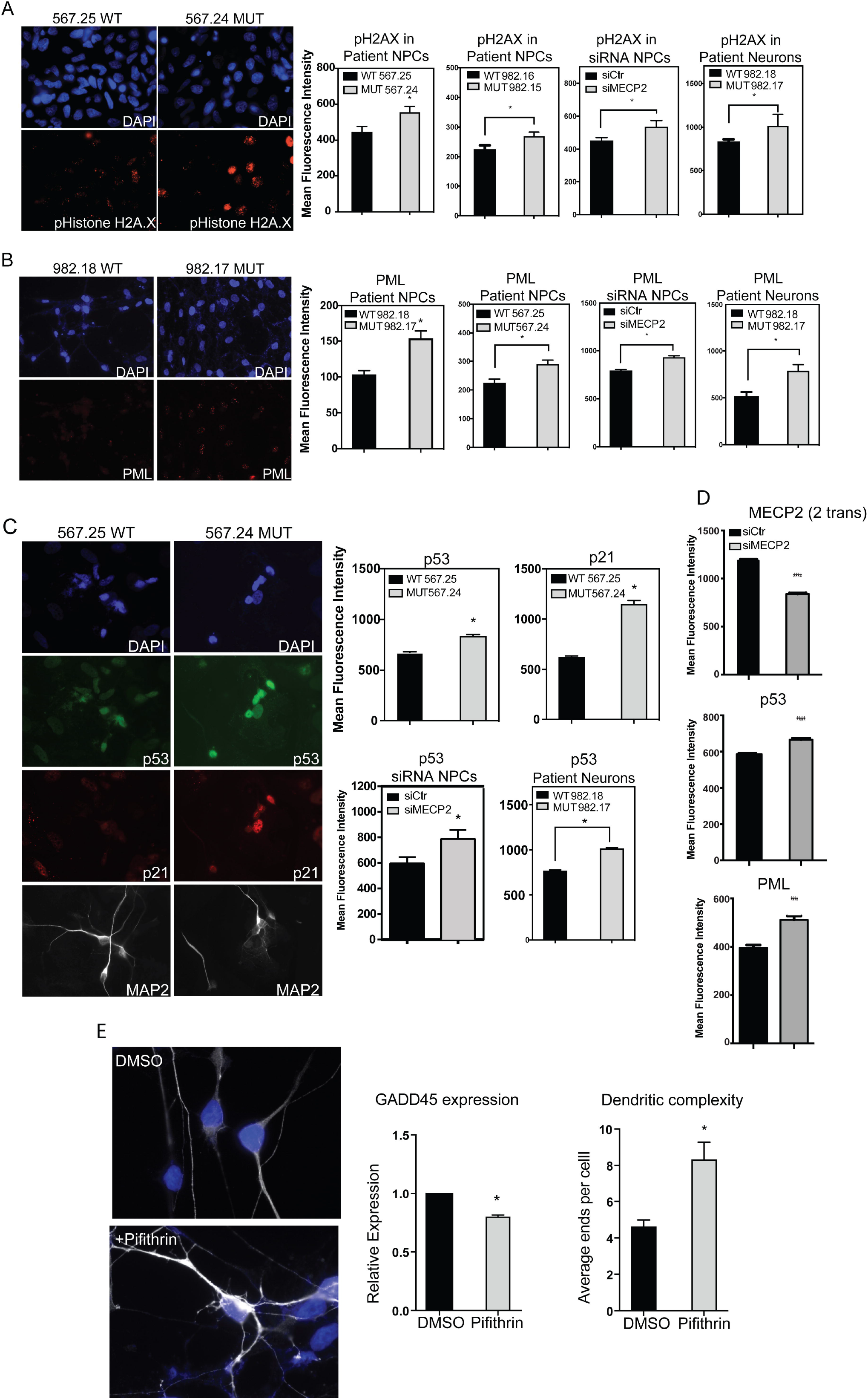
Loss of MECP2 leads to induction of DNA damage and p53. **A**, Immunostaining patient NPCs, NPCs with siRNA against MECP2 and patient neurons in showed a strong increase in H2aX in the absence of MECP2. **B**, Immunostaining patient NPCs, NPCs with siRNA against MECP2 and patient neurons showed a strong increase in PML in the absence of MECP2. **C**, Immunostaining for p53 and p21, a target of p53, showed an increase of these stress markers in MECP2 null neurons. **D**, Immunostaining after siRNA silencing of MECP2 in WT neuronal cultures showed suppression of MECP2 and induction of P53 and PML levels. **E**, Treatment of MECP2-null neurons with DMSO or Pifithrin, followed by immunostaining with antibody for TuJ1 shows a change in dendritic branching and morphology following treatment with Pifithrin. Bottom left, RT-PCR for GADD45, a p53 target gene, showed that Pifithrin reduced p53 activity. Bottom right, Quantification of branching phenotype across three independent experiments showed a strong increase in branching as measured by the number of endpoints. In this figure, all data resulted from at least three independent experiments. *p value<0.05 according to student’s t test. Bar graphs represent mean +/− SEM.

### Blocking induction of P53 can rescue dendritic branching defects due to loss of MECP2

Previous evidence from a murine model of telomere shortening as a result of loss of telomerase complex (TERT) led to defects in dendritic branching, and this effect was strictly dependent on induction of p53^54^. A more recent study also showed that experimentally aging the neural lineage with telomerase inhibition led to neurons with signs of aging, including reduced dendritic branching^55^. Therefore, we posited that inhibition of P53 in MECP2 null neurons could potentially restore appropriate dendritic branching.

To determine whether blocking the action of P53 could improve dendritic branching in MECP2 null interneurons, we took advantage of Pifithrin-α, a potent inhibitor of P53 target gene activation^56^. Treatment of MECP2 null interneurons with Pifithrin-α showed evidence of p53 inhibition as measured by RT-PCR for GADD45^52^, a target gene important for DNA repair (Fig 4E). After 24-48 hours of p53 inhibition by Pifithrin-α, MECP2 null interneurons appeared to adopt an improved neuronal morphology typified by increased physical distinction between the soma and neurites, longer, thinner neurites, as well as increased dendritic branching as shown and quantified in Fig 4E. These data provide evidence that neurons with shortened telomeres due to loss of MECP2 respond by inducing P53 activity, which then inhibits the formation of complex neuronal processes.

## Discussion

Taken together, these data demonstrate that loss of MECP2 leads to clear signs of stress such as H2aX deposition, p53/p21 induction, and initiation of a senescence program, all of which suggest that neurons in Rett Syndrome could be in suboptimal health, leading to neurophysiological defects such as dendritic arborization^13,20^. While one paper suggested that RNAi-mediated silencing of MECP2 could affect the telomeres of mesenchymal cells^37^, decades of work on Rett Syndrome have not uncovered a role for MECP2 in relation to senescence in a wide variety of models such as various transgenic mouse line, human patient post-mortem analyses, *in vitro* human models.

Patients with Rett Syndrome are typically characterized by normal development at birth and subsequent failure to thrive leading to microcephaly and intellectual disability that develops with age. As a result, Rett Syndrome is thought to be caused by experience-dependent loss of neuronal function, which would correlate with data suggesting that MECP2 regulates activity dependent gene expression^10,13,57^. The microcephaly has been proposed to be a function of decreased nuclear size and dendritic arborization of affected neurons^13,20^. Could the induced senescence caused by loss of MECP2 described here underlie patient phenotypes? Several studies have looked at the effects of telomere shortening specifically in the neural lineage and found consistently that upregulation of p53 correlates decreased dendritic arborization^53,54,58^, a phenotype widely described to afflict MECP2 null neurons *in vitro* and *in vivo* (Fig 1).

These results also raise the question of whether senescence could be common to the etiologies of other ID syndromes. The phenotypes described here show a striking similarity to those observed in hiPSCs and neural derivatives made from patients with Immunodeficiency, centromeric region instability, facial anomalies syndrome (ICF) Syndrome^59–61^. Two independent studies showed that ICF patient-derived hiPSCs displayed telomere shortening that was coupled to senescence of somatic derivatives such as fibroblasts. ICF Syndrome only partially overlaps with Rett Syndrome in terms of patient phenotypes, but is caused by mutations in DNMT3B, a *de novo* DNA methyltransferase^62^. These findings together are highly relevant as DNMT3B is a key *de novo* methyl transferase to create methylated DNA (5mC), which is the substrate for Tet oxigenases to create 5-hydroxmethylated DNA (5hmC), which is known to be strongly bound by MECP2^63^. Recently, another study showed that deletion of Tet enzymes, which are critical to generate the 5hmC mark, led to shortened telomeres^64,65^. These studies suggest that DNA hydroxymethylation could be important in the regulation of telomere length. Because MECP2 is a methylated DNA-binding protein, it is tempting to speculate that alterations of methylation lead to defective telomere maintenance.

Another possible interpretation of these data is that instead of a failure to mature, Rett Syndrome neurons instead show aspects of premature aging. The fact that MECP2 null neurons show induction of aging related genes including p53, and induce senescence pathways are consistent with this idea^76^. On the other hand, while Rett patients suffer from a post-natal cognitive decline, and long term survivors show phenotypes associated with Parkinson’s disease^78^, the typical phenotypes presented in young female patients are not consistent with premature aging. Whether the physiological response to loss of MECP2 is truly akin to premature aging or whether patients suffer from the effects that are unrelated to aging is worthy of continued investigation.

Regardless, it is tempting to speculate that treatments that could abrogate the p53 mediated stress response could potentially ameliorate patient outcomes. Pifithrin-**α** has already been shown to be an effective treatment to restore neuronal function in murine models of injury or stroke^79–81^. Significant future effort will be devoted to determining both whether telomere dysfunction is a common trigger for ID Syndromes.

## Acknowledgements

This work was funded by training grants to MO (NIH-Virology and Gene Therapy, UCLA), PL (CIRM, UCLA), CS (CIRM-Bridges, Cal-State-Northridge), DA (HHURP, UCLA). WEL was supported by a Rose Hills Scholar award through the Eli and Edythe Broad Center for Regenerative Medicine. WEL and KP were supported by NIH (P015P01GM099134). This research was also supported by the Allen Distinguished Investigator Program, through The Paul G. Allen Frontiers Group.

## Materials and Methods

### Generation of isogenic Rett Syndrome iPSCs

Two primary fibroblast lines GM17567 (1461A>G in the gene encoding methyl-CpG binding protein 2 (MECP2)), and GM07982 (frameshift mutation, 705delG, in the gene encoding methyl-CpG binding protein 2 (MECP2)), from patients with Rett Syndrome were obtained from Coriell Cell Repositories. 1 × 10^5^ fibroblasts were plated in a gelatin coated well of a 6 well plate in MEF media (DMEM/F12 + 10% FBS). After 8-12 hours, the cells were infected with reprogramming lentivirus that harbors polycystronic human Yamanaka factors (Oct4, Klf4, Sox2, cMyc) in DMEM medium containing 10ug/ml of polybrene and incubated overnight at 37^o^C in 5% CO2 incubator. The following day, the viral media was aspirated, replaced with MEF media and cultured for 3 additional days. Cells were re-plated on the 5th day onto irradiated MEFs in MEF media. On day 6, the culturing media was changed to human ES media containing DMEM/F12 supplemented with L-glutamine, nonessential amino acids (NEAA), penicillin-streptomycin, knockout serum replacement (Invitrogen), and 10 ng/ml basic FGF. Cells were cultured in hiPSC media until iPSC-like colonies were formed. Reprogrammed colonies were further identified by live immunofluorescence staining with TRA-1-81 (Chemicon) then mechanically isolated. Individual colonies were isolated and maintained for at least 2 passages before genotyping analysis. For early passages, the iPSCs were propagated mechanically, whereas collagenase was used for subsequent passaging. hiPSCs were cultured as described previously in accordance with the UCLA ESCRO.

### Generation of teratomas

Generation of teratoma was previously described^82^. Briefly, a single incision was made in the peritoneal cavity of adult SCID mice and the testis was explanted through the incision site. Approximately 60,000 iPSC in a volume of 50 ml 0.5X Matrigel (BD) were transplanted into the testis using a 27-gauge needle. Four to six weeks after surgery, mice were euthanized and the tumors removed for histology. Surgery was performed following Institutional Approval for Appropriate Care and use of Laboratory animals by the UCLA Institutional Animal Care and Use Committee (Chancellor’s Animal Research Committee (ARC)).

### Differentiation in vitro and analysis

Neural specification with neural rosette derivation, neuroprogenitor (NPC) purification, and further differentiation to neurons and glia were performed as described previously^32,83,84^. Relative neuralization efficiency was analyzed by counting the number of neural rosette containing colonies over total number of iPSC colonies. 6-12 35 mm wells were analyzed over four separate experiments. The proliferation efficiency of NPCs was determined by at days 1, 3, and 5 by the total number of cells present in 35mm wells seeded at 200,000 cells on day 0. The cells were detached from the plates using accutase (Millipore) then total number of cells per well analyzed using Z1 Coulter particle counter (Beckman Coulter).

For spontaneous terminal neuronal differentiation by growth factor withdrawal, NPC cultures were subjected to growth factor withdrawal (removal of EGF and FGF) and cultured in basic medium (DMEMF12 + N2 + B27) with three quarter exchange of media every three days. Cells were cultured up to 20 weeks. Neural differentiation efficiency was analyzed four weeks after growth factor withdrawal by counting the number of cells positive for neuronal markers (MAP2 and Tuj1) over the total number of cells visualized by DAPI. NPCs were transfected with DCX-GFP reporter one day prior to differentiation using Lipofectamine 2000 (Invitrogen). Sholl analysis of DCX-GFP positive neuronal neuritis were also measured using ImageJ. All data values were presented as mean +/− SEM. Student’s t-tests were applied to data with two groups. ANOVA analyses were used for comparisons of data with greater than two groups.

For directed differentiation of interneurons, iPSCs were grown on plates coated with matrigel (Corning) until 80% confluency with mTeSR (Stem Cell Technologies). Cells were then treated with DMEM/F12 (GIBCO) containing NEAA (GIBCO), GlutaMAX (GIBCO), bovine serum albumin (Sigma-Aldrich), ß-mercaptoethanol (Sigma-Aldrich), N2 (GIBCO), B27 (GIBCO), SB431542 (10uM; Cayman Chemical), LDN-193189 (100nM; Cayman Chemical) and XAV939 (2uM; Cayman Chemical) later transitioning to the media containing sonic hedgehog (20ng/mL; R&D) and purmorphamine (1uM; Cayman Chemical) as previously described (Maroof et al., 2013). Cells were further differentiated into interneurons with neurobasal medium (GIBCO) containing N2 (GIBCO), B27 (GIBCO), ascorbic acid (Sigma-Aldrich), GlutaMAX (GIBCO), bovine serum albumin (Sigma-Aldrich), ß-mercaptoethanol (Sigma-Aldrich), neurotrophin-3 (10ng/mL; R&D), brain-derived neurotrophic factor (10ng/mL; R&D), and glial cell-derived neurotrophic factor (10ng/mL; R&D).

### Western blot

Cells were lysed on ice with RIPA buffer (Pierce) that contains Halt Protease Inhibitor Cocktail_(Thermo Fisher Scientific) and Halt Phosphatase Inhibitor Cocktail (Thermo Fisher Scientific). The total protein concentration was measured using BCA Protein Assay Kit (Thermo Fisher Scientific) following the manufacturer’s protocol. The lysates containing the equal amount of total protein were mixed with NuPAGE sample buffer (Invitrogen) with 5% mercaptoethanol and denatured at 95 °C for 10 min. Supernatant was electrophoresed onto NuPAGE 4-12% Bis-Tris Protein Gels (Invitrogen) using MOPS running buffer (Invitrogen). Gels were then electroblotted using semi-dry transfer apparatus onto a membrane. The membrane was blocked with 5% non-fat milk for 1 hr and incubated overnight with primary antibodies at 4°C. The next day the membrane was washed and incubated with HRP-conjugated secondary antibodies for 1 hr at room temperature. The membrane was then incubated with ECL Western Blotting Substrate and developed.

### Immunofluorescence and image quantification

Cells on coverslips were washed with PBS, fixed in 4% paraformaldehyde for 15 min at room temperature, blocked for 1 hr at room temperature with 10% serum and 0.1% Triton-X-100, then incubated overnight at 4 °C with primary antibodies. Frozen tissue sections were thawed to room temperature, fixed in 4% paraformaldehyde for 15 min at room temperature, permeabilized with 0.2% Triton-X-100 for 15 min at room temperature and blocked with 10% serum at 4 °C overnight, followed by incubation with primary antibodies at 4 °C for 16-24 hr. Following primary antibody incubation, the coverslips were incubated with Alexa Fluor (Invitrogen) or Jackson immunoresearch secondary antibodies at room temperature for 1 hr. Cells were counterstained with DAPI and mounted in Prolong Gold (ThermoFisher). Antibodies used include the following: mouse anti-OCT3/4 (1:100, Santa Cruz Biotechnology Inc.), rabbit anti-SOX2 (1:300, Cell Signaling Technology), rabbit anti-Nanog (1:100, Cell Signaling Technology), mouse anti-Tra-1-81 (1:250, Chemicon), mouse anti-NESTIN (1:1000, Neuromics), chicken anti-MAP2 (1:2000, Biolegend), chicken anti-GFAP (1:2000, Abcam), rabbit anti-Tubulin β3 (1:500, Covance), mouse anti-p53 (1:500, Cell Signaling), rabbit anti-p21 (1:250, Santa Cruz), mouse anti-PML (1:100, Santa Cruz), mouse anti-phospho-Histone H2A.X (1:2000, EMD Millipore), rabbit anti-5hmc (1:100, Active Motif), rabbit anti MECP2 (1:1000, Diagenode), rabbit anti Foxg1 (1:1000, Abcam), and mouse anti NKX2.1 (1:300, Novocastra). Secondary antibodies conjugated with Alexa 488, 568, 594, 647 (1:500, Life Technologies) were used. Imaging was performed on Zeiss Axio Imager A1 or Zeiss LSM780 confocal microscope using a 40X or 63X objective on randomly selected cells. Mean intensity or a number of foci were quantified using ImageJ (http://rsb.info.nih.gov/ij/). At least 100 cells per condition were used for each independent experiment.

### RT-qPCR

RNA from cultured cells was collected using the RNeasy Mini Kit from Qiagen according to the manufacturer’s instructions. The concentration and purity of RNA were measured using nanodrop spectrophotometers (Thermo Scientific). RNA with an A260/A280 ratio in between 1.8 and 2.0 as well as an A260/A230 ratio in between 2.0 and 2.2 was used. RNA was then reverse transcribed using the Super Script III First-Strand cDNA Synthesis kit with Random Hexamers (Invitrogen) according to the manufacturer’s instructions. Quantitative PCR was performed using SYBR Green master mix (Roche). Primers were used at a final concentration of 1 uM. Reactions were performed in duplicate and duplicate CT values were averaged and then used for standard ΔΔCT analysis. Expression levels were normalized to beta actin.

### Data collection and statistical analysis

All the experimental data (RT-qPCR, qPCR assay for telomere length, immunostaining, ß-Galactosidase Senescence Assay) were presented as mean +/− SEM based on at least three biological replicates from independent experiments. Student’s t tests were applied to data with two groups. ANOVA analyses were used for comparisons of data with greater than two groups. A p value < 0.05 was considered as statistically significant.

### siRNA gene silencing

All knockdown experiments were performed using trilencer siRNAs (from OriGene Technologies) and RNAimax (ThermoFisher) in Opti-MEM media (ThermoFisher). Trilencers were used at a concentration of 20 nM. Transfection media was prepared and then 500,000 cells were plated on top of the transfection media in 6-well plates. The medium was changed to normal NPC media the next day and cells were collected for analysis at the time points indicated.

### ß-Galactosidase Senescence Assay

ß-Galactosidase Senescence Assay was performed using the Senescence β-Galactosidase Staining Kit from Cell Signaling according to manufacturer’s instructions. Briefly, the cells were fixed on coverslips, incubated with X-gal overnight at 37°C, then mounted on glass slides and imaged using a brightfield microscope. The number of blue cells and number of total cells were quantified using the Cell Counter plugin in ImageJ.

### Quantification of Dendritic Arborization

Neuronal cultures were immunostained for Tuj1 in order to identify mature neurons and visualize entire cells. The stained cells were then imaged at 20x and dendritic arbors of individual cells were traced using the Simple Neurite Tracer plugin for ImageJ. The number of process ends per cell were counted using the Cell Counter plugin for ImageJ. Only mature neurons—identifiable by their thin processes and condensed somas—were used for analysis. The number of process ends per cell are presented as mean ends per cell +/− SEM. Means were compared using the Student’s T-Test for data with two groups. A p value <0.05 was used as the cutoff for significance.

### RNA expression profiling

RNA purification was performed with Qiagen RNAeasy kit following the manufacturer’s instruction. Libraries were prepared according to the manufacturers guidelines using The TruSeq V2 kit (Illumina). Microarray profiling was performed with Affymetrix Human HG-U133 2.0 Plus arrays as described^85^. For RNA sequencing, the datasets were mapped with RASER and HISAT2. The reads were counted per gene. Genes were defined by the exon union from the hg19 ensembl annotations. The function of DESeq in DESeq2 package was used to first normalize the gene read counts data and then identified the differentially expressed genes. The MA plot was generated with the function of plotMA in DESeq2 package. Q-value of 0.05 is regarded as the stringent cutoff of calling DEGs while p-value less than 0.05 is regarded as the low stringency cutoff. For the meta-chromosome plot of DEGs, all the chromosomes (except chromosome Y) were first divided equally into 20bins with different length, and then the number of DEGs in each bin was counted. GO analysis was performed using DAVID.

---

**Figure S1.** Validation of disease in a dish model for Rett Syndrome **A**, Teratoma assay was performed to establish pluripotency of hiPSCs made from Rett patient fibroblasts. The resulting tumors each showed evidence of differentiation towards all three embryonic germ layers. **B**, NPCs were produced from isogenic hiPSCs of Rett patient, and assessed by western blot to validate loss of MECP2 and specificity of antibody. Top panel shows that the antibody only recognizes MECP2. Bottom panel shows that in NPCs from both patients, individual clones either express or lack MECP2. **C**, The ability of hiPSCs to generate NPCs was assayed in Rosette formation assay. Lack of MECP2 did not affect rosette formation across multiple lines from both patients. N=4 independent experiments. *p value < 0.05 according to student’s t test (for patient R567) or ANOVA (for patient R982). Bar graphs represent mean +/− SEM. **D**, Growth curves show that loss of MECP2 does not affect proliferation of NPCs made from either patient. **E**, 3 weeks of growth factor withdrawal drives NPCs to differentiate into neurons and glia as measured here by immunostaining for MAP2/Tuj1 or S100/GFAP in patient 567 derived cultures. There was no consistent difference in differentiation potential across lines from either patient. N=2 independent experiments. Bar graphs represent mean +/− SEM. **F**, Patient 982 derived cultures also do not show dramatic differences in the presence of neurons or astrocytes as measured by MAP2 and S100. N=3 independent experiments. Bar graphs represent mean +/− SEM.

**Figure S2.** RNA-seq analysis to determine the relative ration of WT versus MUT transcripts of MECP2 in Rett patient derived lines. **A and B**, immunostaining to demonstrate the efficiency of directed differentiation towards neural progenitors (A) and then onto interneurons (B) with markers typical of each stage. **C**, Detection of WT and MUT transcripts from each of the lines indicated demonstrated a clear bias towards individual alleles in each patient derived line. This analysis indicates XCI status for each allele, and demonstrates that XCI status is unchanged, even after differentiation to neurons.

**Figure S3.** Silencing MECP2 by siRNA MECP2 was downregulated by RNA interference, quantified by RT-PCR (left), for protein by western blot (middle), and as demonstrated by immunostaining for MECP2 (right). N=3 independent experiments. *p value < 0.05 according to student’s t test. Bar graphs represent mean +/− SEM.

